# Effects of menopause and high fat diet on metabolic outcomes in a mouse model of Alzheimer’s disease

**DOI:** 10.1101/2023.11.21.568069

**Authors:** Charly Abi-Ghanem, Abigail E. Salinero, David Riccio, Richard D. Kelly, Krystyna A. Rybka, Olivia J. Gannon, David Kordit, Nyi-Rein Kyaw, Febronia Mansour, Kasey M. Belanger, Christina A. Thrasher, Matthew Wang, Emily Groom, Rachel M. Smith, Heddwen L. Brooks, Lisa S. Robison, Damian G. Zuloaga, Kristen L. Zuloaga

**Affiliations:** Department of Neuroscience & Experimental Therapeutics, Albany Medical College, 47 New Scotland Avenue; MC-136, Albany, NY 12208, USA; Department of Psychology and Center for Neuroscience Research, State University of New York at Albany, 1400 Washington Ave, Biology 325, Albany, NY 12222, USA; Department of Physiology, Tulane University School of Medicine, New Orleans, LA 70112, USA; Department of Psychology and Neuroscience, Nova Southeastern University. 3300 S University Drive, Fort Lauderdale, FL 33328, USA

**Author notes:** Corresponding author: Kristen L. Zuloaga, PhD, Associate Professor, Department of Neuroscience & Experimental Therapeutics, Albany Medical College, 47 New Scotland Avenue; MC-136 Albany, NY, USA 12208.

**Keywords:** Metabolic disease, Alzheimer’s disease, Menopause, Hypothalamus, Obesity, Glucose metabolism

## Abstract

About two-thirds of those with Alzheimer’s disease (AD) are women, most of whom are post-menopausal. Menopause accelerates the risk for dementia by increasing the risk for metabolic, cardiovascular, and cerebrovascular diseases. Mid-life metabolic disease (e.g. obesity, diabetes, or prediabetes) is a well-known risk factor for dementia. A high fat diet can lead to poor metabolic health in both humans and rodents. The goal of this study was to determine the effects of menopause and high fat diet on metabolic outcomes in the App^NL-F^ knock-in mouse model of Alzheimer’s disease. To model menopause, we used an accelerated ovarian failure model (4-vinylcyclohexene diepoxide, VCD). This ovary-intact model is more clinically relevant than an ovariectomy model, as mice go through a perimenopausal period. At 3 months of age, App^NL-F^ mice were administered VCD or vehicle (oil) and then placed on either a control diet (10% fat) or a high fat diet (HF; 60% fat) and maintained on the diets until 10 months of age. Menopause led to metabolic impairment (weight gain and glucose intolerance) and further exacerbated obesity in response to a high fat diet. Menopause had independent effects on some serum metabolic health biomarkers (insulin) and synergic effects with HF diet on other markers (glucagon). Some metabolic effects of menopause may be centrally mediated, as menopause altered the expression of hypothalamic genes related to energy balance and increased microgliosis in the lateral hypothalamic nucleus. This work highlights the need to model endocrine aging in animal models of dementia and will contribute to further understanding the interaction between menopause and metabolic health in the context of AD.

**Highlights:**

- In a mouse model of AD, menopause, modeled by accelerated ovarian failure, leads to metabolic impairment.
- Menopause has independent effects on some serum metabolic health biomarkers (insulin) and synergic effects with HF diet on other markers (glucagon).
- Menopause alters the expression of hypothalamic energy balance related genes.
- Menopause leads to increased microgliosis in the lateral hypothalamic nucleus.

## Introduction

Worldwide, it is estimated that about 22% of people over 50 years of age suffer from Alzheimer’s disease (AD) at various clinical stages (1). Currently, approximately 6 million Americans are living with AD. This number is rapidly increasing, and it is estimated that in 2060, about 14 million Americans will have AD (2). AD neuropathology, such as amyloid plaques, neurofibrillary tangles, and neuroinflammation, begins accumulating in mid-life, decades before cognitive impairment and diagnosis. Since some of the most striking symptoms associated with AD are related to cognitive decline, the most commonly studied brain regions are those that are implicated in cognitive function. However, metabolic dysfunction is another symptom of AD that has recently been gaining attention. The hypothalamus, responsible for the regulation of several homeostatic functions, is affected by AD pathology, resulting in the dysregulation of metabolism, sleep, and neuroendocrine alterations in AD patients (3, 4). Previously, we and others have reported that metabolic disturbances are associated with pathological changes to the hypothalamus in rodent models of AD (3, 5-8).

Not only does AD contribute to hypothalamic and metabolic disturbances, but conversely, metabolic disturbances are a driver of AD pathology (3). Metabolic disease (including obesity and diabetes or prediabetes), especially in mid-life, is a major risk factor for developing AD (4, 9-11). A recent study showed that in individuals older than 60 years of age, metabolic disease was associated with a greater than 11-fold higher risk of AD compared to metabolically healthy individuals (11). Additionally, up to 80% of AD patients suffer from impaired glucose metabolism (12) and insulin resistance has been reported in the brain of AD patients (13, 14). Animal studies, including our own, are in line with these clinical observations. Inducing metabolic disease via consumption of a high fat (HF) diet has been shown to worsen AD pathology and cognitive deficits in animal models (5, 15-19). Previously, we reported that consumption of a HF diet from ∼3-7 months of age in 3xTg-AD mice resulted in a wider array of cognitive deficits (20) as well as more severe weight gain, glucose intolerance, and hypothalamic inflammation in females compared to males (5). These results not only highlight the association between metabolic impairment and AD, but also shed light on underlying sex differences, with a greater vulnerability of females.

Female sex is a major risk factor for AD, with women constituting 2/3 of all those suffering from the disease (21, 22). Since the greatest risk factor for AD is advanced age, most women with AD are post-menopausal. Menopause is a mid-life endocrine transition during which women experience a loss of ovarian hormones, leading to increased risk for cognitive decline and dementia (23-27). This is in part due to the loss of the neuroprotective effects of estrogens, as well as the exacerbation of dementia risk factors (24, 25, 27, 28), as menopause has been associated with increased weight gain and visceral fat accumulation, increased rates of type 2 diabetes, and altered brain glucose metabolism (29-32).

In this study we sought to determine the effect of menopause on metabolic disease in the context of AD. Using a knock-in mouse model of AD, App^NL-F^ mice (33), we induced metabolic disease using a HF diet and modeled menopause using an accelerated ovarian failure model. This ovary-intact accelerated ovarian failure model of menopause is more clinically relevant than an ovariectomy model, as the mice go through a perimenopausal period (34). We found that both menopause and HF diet independently or synergistically had negative effects on several metabolic outcomes, potentially through modulation of hypothalamic gene expression and neuroinflammation.

## Methods

### Animals and experimental design

All experiments were approved by the Albany Medical College Animal Care and Use Committee and in compliance with the ARRIVE guidelines. App^NL-F^ (App^tm2.1Tcs^/App^tm2.1Tcs^) mice (33) were acquired congenic on a C57BL/6J background from Dr. John Cirrito at Washington University following MTA from Riken, Japan. Mice were bred in house and housed (3-4 per cage) at ∼21°C, 30-70% humidity, with a 12 h light/dark cycle. At three months of age, cages of female App^NL-F^ (App^tm2.1Tcs^/App^tm2.1Tcs^) mice were randomized to treatment groups and placed on either a high fat (HF) diet (60 kcal% fat, D12492, Research Diets, New Brunswick, NJ, USA) or a low fat (LF) diet (10 kcal% fat, D12450B, Research Diets, New Brunswick, NJ, USA). To model menopause, we induced accelerated ovarian failure as described previously (35). Starting at diet onset (3 months of age), mice received once daily i.p. injections of 4-vinylcyclohexene diepoxide (VCD, 160 mg/kg) for 20 consecutive days. Control mice on each diet received vehicle (sesame oil) injections. Cyclicity was assessed using vaginal cytology starting 2 months after the 1st injection (mice ∼ 5.5 months old), and mice that remained in diestrus for 10 consecutive days were declared acyclic/menopausal. Six months after diet onset, mice received a glucose tolerance test (GTT) to evaluate their diabetic status, as described previously (5, 36, 37). One month later (at 10 months old), mice were euthanized by deep anesthesia followed by cardiac puncture to collect blood, then perfused with ice cold heparinized saline. Brains were collected and bisected, with one hemisphere randomly selected to undergo post-fixation for use with immunofluorescent labeling, and the other hemisphere regionally dissected, and flash frozen for qPCR. Visceral, subcutaneous, and brown fat weights were collected at endpoint. A timeline of the experiment is provided in **Figure 1**.

**Figure 1:**
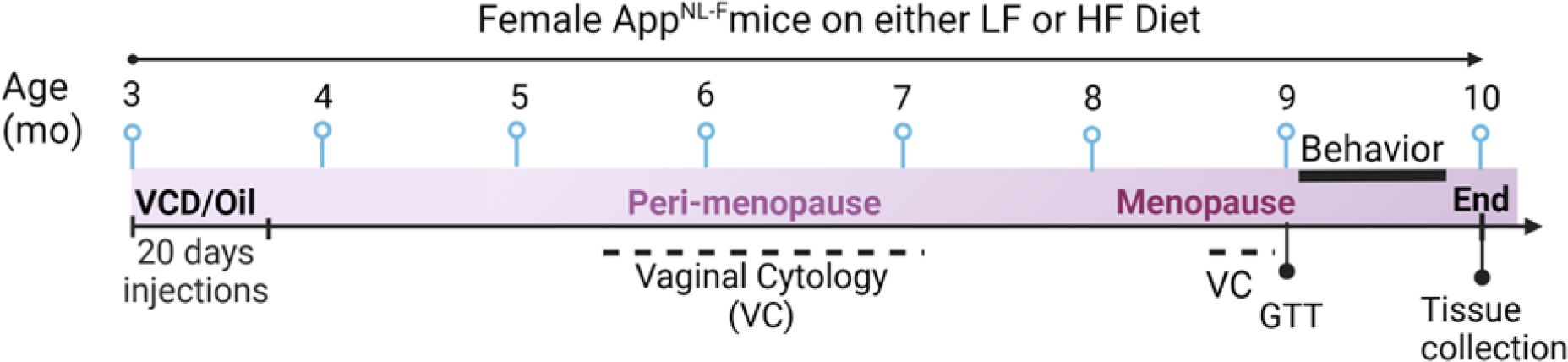
Experimental Timeline. Alzheimer’s disease is modeled using female App^NL-F^ mice. To model menopause, we used a chemically-induced accelerated follicular atresia model by injecting mice for 20 consecutive days with 4-vinylcyclohexene diepoxide (VCD). Control mice received oil (vehicle) injections. At 3 months of age, mice were put on either a high fat (HF, 60% fat) or low fat (control, LF, 10% fat) diet and oil or VCD injections started. The Estrus cycle was monitored using vaginal cytology (VC) starting 60 days after the first injection. Mice were declared menopausal when they had 10 consecutive days in the diestrus phase. At about 9 mo of age, mice were confirmed menopausal or cycling depending on their experimental group, then their prediabetic state was evaluated using glucose tolerance test (GTT). One month later, mice were euthanized for end point measures. Figure created with BioRender.

### Serum diabetes markers

During euthanasia, blood was collected via cardiac puncture and left to coagulate then spun at 1500×g for 10 min at 4 °C to collect serum. Serum samples were aliquoted and stored at -80°C until assayed. Diabetes-associated markers were assessed using the Bio-Plex Pro Mouse Diabetes 8-Plex Assay (171F7001M; Bio-Rad, Carlsbad, California) according to the manufacturer’s instructions.

### Quantitative polymerase chain reaction (qPCR)

RNA was extracted from flash frozen hypothalamic tissue using Qiagen RNeasy plus mini kit according to the manufacturer’s instructions (74131 Qiagen USA). Next, 240ng of RNA were converted to cDNA using High-Capacity cDNA Reverse Transcription Kit with RNase Inhibitor (4374967 Thermo Fisher Scientific) according to the manufacturer’s instructions. Quantitative PCR was performed in triplicates on 10ng of cDNA in a 10μl reaction using TaqMan probe technology on a Bio-Rad CFX-384 real time system. Taqman assays (Thermo Fisher Scientific) used were: Agrp (Mm00475829_g1), Npy (Mm01410146_m1), Pomc (Mm00435874_m1), Lepr (Mm00440181_m1), Mc4r (Mm00457483_s1). The housekeeping genes used were Rpl13a (Mm01612986_gH) and Rps17 (Mm01314921_g1). Bio-Rad CFX Maestro 1.1 software was used to analyze the data. The relative expression levels of genes of interest were calculated using the ΔΔCq method relative to either of the housekeeping genes using the females on a LF diet with oil injections as reference group.

### Microglia analysis

Hemispheres used for immunofluorescent labeling were fixed overnight in a 4% paraformaldehyde (PFA) solution, cryoprotected in 30% sucrose, then frozen in optimal cutting temperature (O.C.T.) solution (23-730-571, Thermo Fisher Scientific) and stored at -80°C until further processing. Tissue sections (35μm thickness) were obtained using a cryostat (Cm1950, Leica) and stored at 4 °C in cryopreserve. Immunofluorescent labeling was performed as previously described (5, 20, 38). Briefly, slices were permeabilized and blocked for one hour at room temperature using 0.3% Triton X-100 in PBS (TPBS) with 5% donkey serum solution. Primary antibody of goat anti-Iba-1 (1:1000; PA5-18039, lot #TI2638761, Thermo Fisher Scientific) was applied overnight at 4°C in blocking solution. Alexa Fluor 647 donkey anti-goat (1:300; 705-605-147, Jackson ImmunoResearch) was added in blocking buffer for 2h at room temperature. Slices were counter stained with DAPI and washed 3 times before mounting between slides and coversliped using ProLong™ Gold Antifade Mountant (P36930 Thermo Fisher scientific). Images of brain slices were obtained at 10x magnification using the Axio Observer Fluorescent Microscope (Carl Zeiss Microscopy, Oberkochen, Germany). Regions of interest (ROIs) were drawn around the arcuate nucleus (ARC), dorsomedial hypothalamus (DMH), ventromedial hypothalamus (VMH), lateral hypothalamus area (LH), and paraventricular nucleus (PVN) of every 8th section using ImageJ software (NIH, Bethesda, MD, USA) to quantify the average area covered (%). All measurements were made by an experimenter who was blinded to the treatment group.

### Open field testing

To test for locomotor activity, mice were placed in a square arena (45x45cm) and allowed to explore freely for 10 min, then removed and placed in a “recovery cage” so as not to expose them to naïve cage mates. Videos were analyzed using ANY-maze software (Stoelting, Wood Dale, IL). Total distance traveled (m) was measured as a proxy for locomotor activity.

### Statistical analysis

Statistical analyses were completed using GraphPad Prism (GraphPad Software v10, San Diego, CA, USA). Statistical outliers were removed following identification using Grubbs’ test with alpha set at 0.05. Data were analyzed using 2-way ANOVAs followed by Fisher’s LSD post hoc test, except for measures tracked over time (GTT) that used 3-way repeated measures ANOVAs. Correlations were run for all animals and separately for each group using Pearson’s correlations. Statistical significance was set at p<0.05. Data are expressed as mean+SEM.

## Results

### Menopause and HF diet cause weight gain and glucose intolerance in App^NL-F^ mice

Metabolic deficits have been observed in AD patients and postmenopausal women (3, 24). In a mouse model of AD, we assessed differences in the effect of HF diet and menopause on several metabolic outcomes such as body weight, adiposity, and glucose tolerance. Unsurprisingly, females on a HF diet weighed significantly more than those on the LF diet (**Figure 2 A**, main effect of diet p<0.0001). We found a main effect of menopause in increasing body weight (**Figure 2A**, main effect of menopause p=0.006), with VCD injected (menopausal) mice weighing more than those injected with oil. We sought to determine whether the increase in body weight was due to differences in adiposity. HF diet increased visceral fat (**Figure 2B**, main effect of diet p<0.0001), subcutaneous fat (**Figure 2C**, main effect of diet p<0.0001), and brown fat weights (**Figure 2D**, main effect of diet p=0.0035). No effect of menopause was observed for fat accumulation.

**Figure 2:**
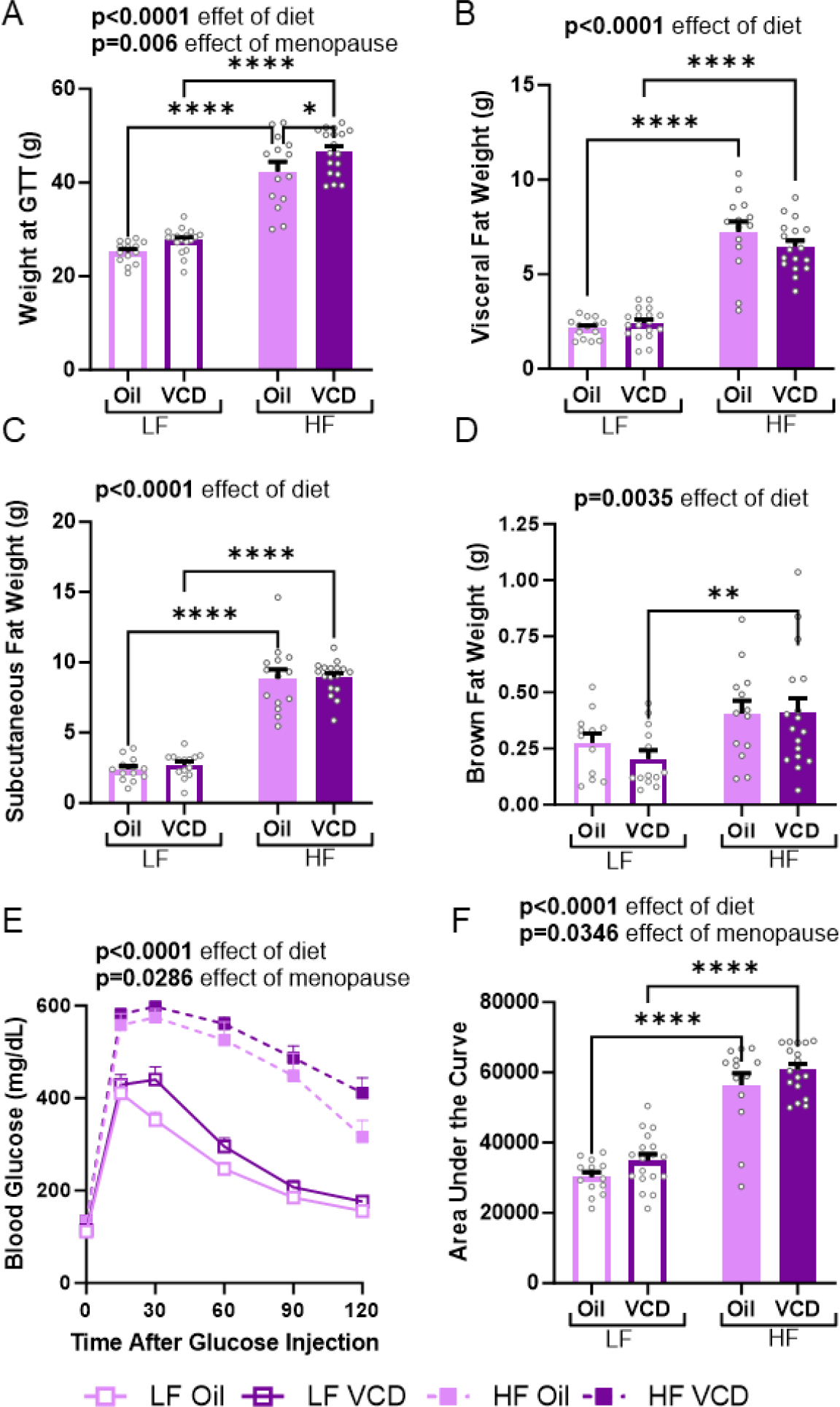
Metabolic Effects of HF Diet and Menopause in Female APP^NL-F^ Mice. Metabolic status was assessed using body weight, adiposity, and glucose intolerance as outcome measures. Body weight was measured at the time of glucose tolerance test (GTT, ∼9mo; **A**). Adiposity was evaluated by measuring the wet weights of several fat tissue types at endpoint, including visceral fat (**B**), subcutaneous fat (**C**), and brown fat (**D**). Prediabetic status was evaluated using GTT. (**E**) Blood glucose levels (mg/dL) before (fasting) and up to 2 hours after injection of a glucose bolus (2 g/kg in saline i.p.). (**F**) Area under the curve for blood glucose during GTT. A 2way ANOVA was used to compare the effect of diet and menopause on each variable. For the GTT (**E**), a 3way repeated measures ANOVA was used. Main effects are indicated above the graphs. *** p<0.001, **** p<0.0001. n=14-19 mice/group. High fat (HF), low fat (LF), 4-vinylcyclohexene diepoxide (VCD, menopause model).

Additionally, we evaluated the prediabetic status of mice by performing a GTT 6 months after diet onset (∼9mo of age). When examining glucose clearance over time, there was a main effect of HF diet to impair glucose clearance (**Figure 2E**, main effect of diet p<0.0001). This diet effect translated into a larger area under the curve (AUC, worse glucose intolerance) in mice on a HF diet compared to those on a LF diet (**Figure 2F**, main effect of diet p<0.0001). Additionally, we observed a main effect of menopause in worsening glucose intolerance, independent of diet (**Figure 2E**, main effect of menopause p=0.0286). This was supported by a larger AUC for VCD injected mice (**Figure 2F**, main effect of menopause p=0.0346). Taken together, these results show that menopause, on its own or in addition to HF diet, led to metabolic impairment in App^NL-F^ mice.

### Effects of menopause and HF diet on serum levels of diabetes-associated markers

Since HF diet and menopause had effects on metabolic outcomes, we sought to assess serum levels of several diabetes-associated markers. Leptin is a hormone that is released by adipose tissue to limit food intake (39). As expected, HF diet resulted in hyperleptinemia, with leptin levels in HF fed mice reaching ∼3-fold higher than those on a LF diet (**Figure 3A**, main effect of diet p<0.0001), but menopause had no effect on leptin levels. No group differences were observed in ghrelin levels, a hormone that stimulates feeding behavior (39) (**Figure 3 B**). Next, we examined hormones that control blood glucose levels, including insulin (decreases blood glucose) and glucagon (increases blood glucose). As expected, HF diet increased insulin levels (**Figure 3C**, main effect of diet p=0.0361). Additionally, we observed a trend in menopause effects (p=0.077), which was driven by a significant decrease in insulin levels in LF VCD-injected mice compared to LF oil-injected controls (p=0.0311). HF diet led to decreased glucagon levels (**Figure 3D**, main effect of diet p=0.0025). Post-hoc tests revealed that the decrease in glucagon was mediated by a reduction in the HF menopausal mice (LF VCD vs HF VCD p=0.007). As expected, HF diet also increased levels of resistin (a hormone that suppresses insulin function) (**Figure 3E**, main effect of diet p<0.0001) and plasminogen activator Inhibitor-1 (PAI-1, a biomarker of fibrinolysis that has been shown to increase with age, obesity, and insulin resistance (40)) (**Figure 3F**, main effect of diet p<0.0001). Menopause did not alter either of these markers. No differences were observed between groups for levels of gastric inhibitory polypeptide (GIP; **Figure 3G**) and glucagon-like-peptide 1 (GLP-1, **Figure 3H**); these hormones have opposing actions on glucagon secretion, respectively enhancing, and repressing it (41). Taken together, these results show that menopause can have independent (insulin) or synergistic (glucagon) effects with HF diet on serum levels of metabolic hormones in App^NL-F^ female mice.

**Figure 3:**
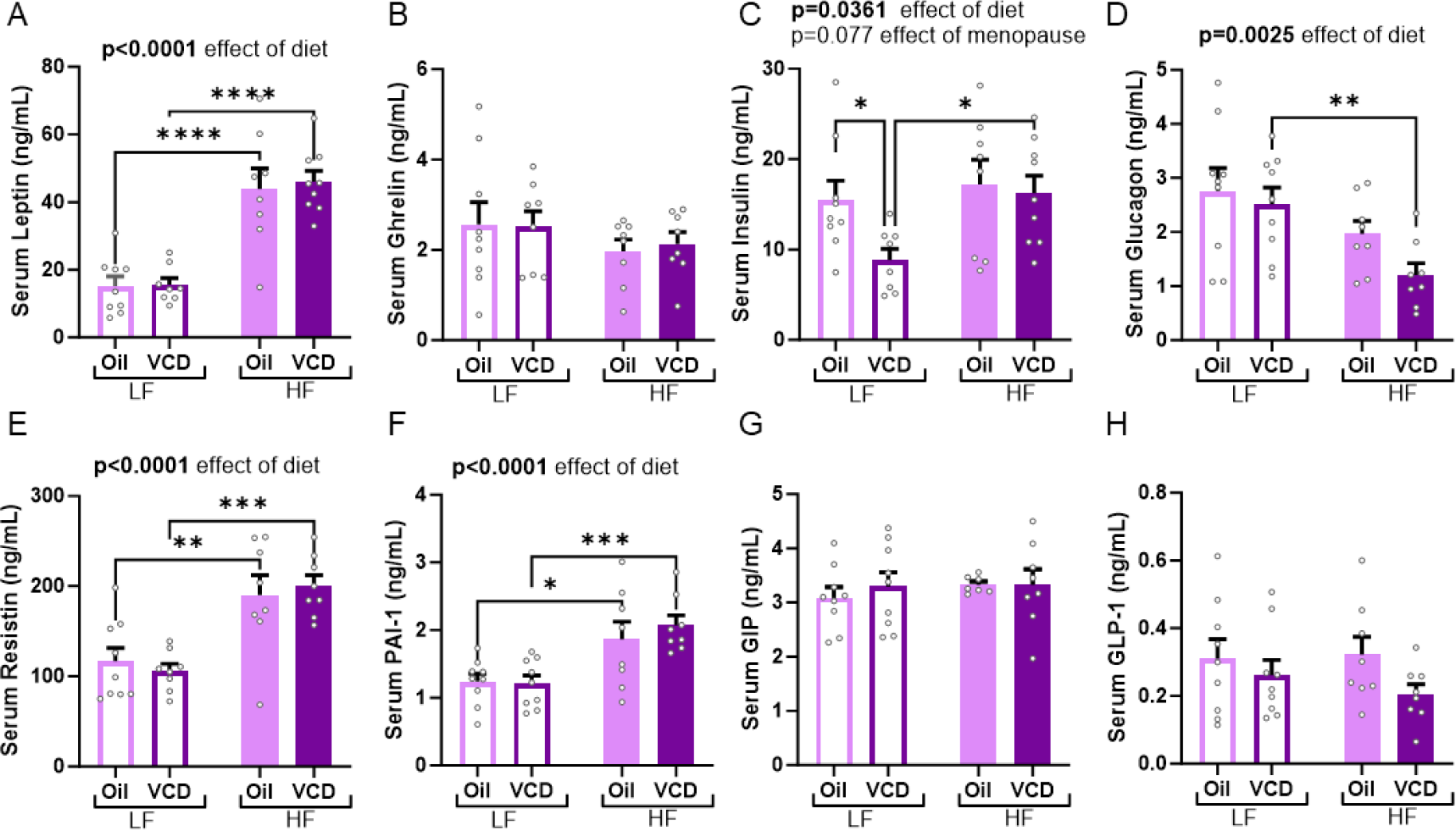
Effects of HF Diet and Menopause on Serum Levels of Diabetes-associated Markers. At end point (∼7mo after diet onset, mice ∼10 mo), blood was collected via cardiac puncture and serum was extracted and stored at -80°C. Diabetes associated markers were later measured using the Bio-Plex Pro Mouse Diabetes 8-Plex Assay. Graphs represent serum levels (ng/mL) of leptin (**A**), ghrelin (**B**), insulin (**C**), glucagon (**D**), resistin (**E**), plasminogen activator inhibitor-1 (PAI-1; **F**), gastric inhibitory polypeptide (GIP; **G**) and glucagon-like-peptide 1 (GLP-1; **H**). A 2way ANOVA was used to compare the effect of diet and menopause on each marker. Main effects are indicated above the graphs. * p<0.05, **p<0.01, *** p<0.001, **** p<0.000.1 n=8-9 mice/group. High fat (HF), low fat (LF), 4-vinylcyclohexene diepoxide (VCD, menopause model).

### Interaction between menopause and HF diet on hypothalamic markers of energy balance

Hypothalamic dysfunction, which can lead to disturbances in feeding behavior and metabolic function, is well known to occur in AD (3). Therefore, we measured the expression of several hypothalamic genes involved in regulating metabolic function and feeding behavior. We did not observe any changes in the expression of the fast-acting orexigenic neuropeptide Y (Npy, **Figure 4A**). However, menopause altered levels of the delayed, longer-acting orexigenic agouti-related peptide (Agrp, **Figure 4B**) (main effect of menopause p=0.037), with a decrease in Agrp mRNA expression specifically in menopausal females on a LF diet (LF oil vs. LF VCD p=0.0054). This resulted in a menopause by diet interaction (p=0.040). Next, we examined the expression of melanocortin 4 receptor (Mc4r), because Mc4r deficiency is linked to hyperphagia (42, 43). HF diet trended towards decreasing Mc4r (main effect of diet p=0.062); conversely, menopause trended towards increasing Mc4r (**Figure 4C**, main effect of menopause p=0.053). No differences were observed between groups in expression levels of leptin receptor (Lepr, **Figure 4D**) or pro-opiomelanocortin (Pomc, **Figure 4E**), which play roles in energy balance.

**Figure 4:**
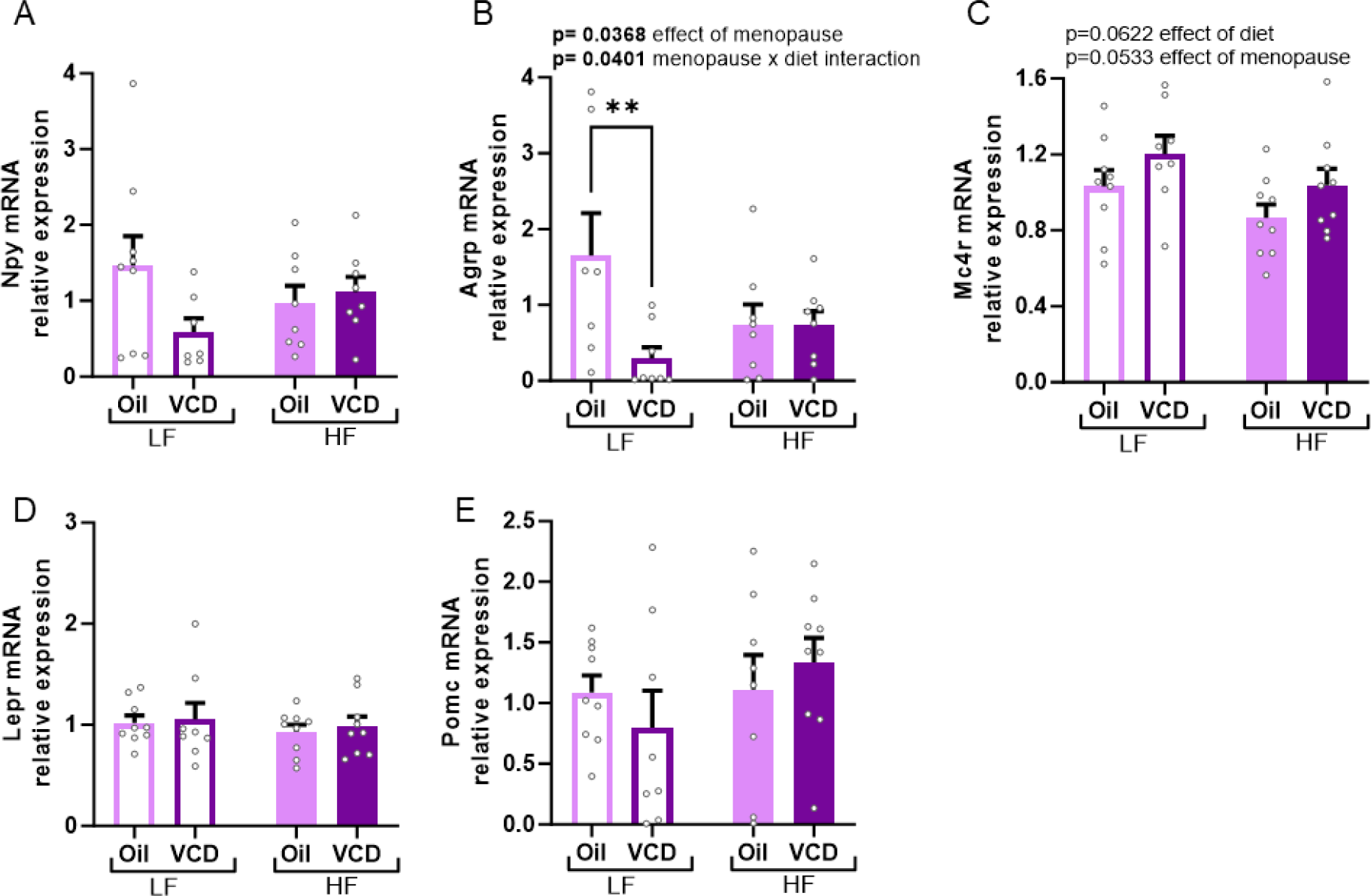
Interaction Between Menopause and HF Diet on the Expression of Hypothalamic Markers. Hypothalami were dissected, flash frozen and then RNA was extracted. qPCR was used to measure the expression levels of mRNA of genes coding for several markers related to food consumption and energy balance. Graphs represent relative expression levels (fold change compared to LF oil mice) of Npy (**A**), Agrp (**B)**, Mc4r (**C**), Lepr (**D**), Pomc (**E**). A 2way ANOVA was used to compare the effect of diet and menopause on each marker. Main effects are indicated above the graphs. ** p<0.01, # p<0.05 main effect of menopause, n=7-9 mice/group. High fat (HF), low fat (LF), 4-vinylcyclohexene diepoxide (VCD, menopause model).

We examined the association between these hypothalamic genes, metabolic measures, and serum levels of diabetes-associated markers (**Sup Figure 1**). We observed different associations in oil-injected controls versus VCD-injected menopausal mice between hypothalamic gene expression and metabolic markers (**Sup Figure 1B**) and serum levels of diabetes associated hormones (**Sup Figure 1C**). For example, in menopausal mice, but not in controls, Mc4r expression was significantly associated with several metabolic measures, as well as insulin and leptin levels. On the other hand, ghrelin levels were negatively associated with LepR expression in control but not menopausal mice. These results indicate a potential shift in metabolic control post-menopause, and that some menopause-induced metabolic changes might in part result from central effects on hypothalamic gene expression.

### Menopause increases microgliosis in some hypothalamic regions

Microglia activation is a prominent feature of AD (44). Additionally, in both humans and rodent models, obesity is associated with increased gliosis notably in the hypothalamus (45-47). Immunolabeling was performed for Iba1 to assess microgliosis in several hypothalamic nuclei (**Figure 5**). In the lateral hypothalamus (LH, **Figure 5A**), menopause increased Iba1+ density regardles of diet (main effect of menopause p=0.0404). There were no signficant effects of HF diet or menopause in the dorsomedial hypothalamus (DMH), the ventromedial hypothalamus (VMH), the arcuate nucleus (ARC) or the paraventricular nucleus (PVN; **Figure 5B-E**). These results show that the lateral hypothalamus is particulary vulnerable to increased microgliosis in response to menopause.

**Figure 5:**
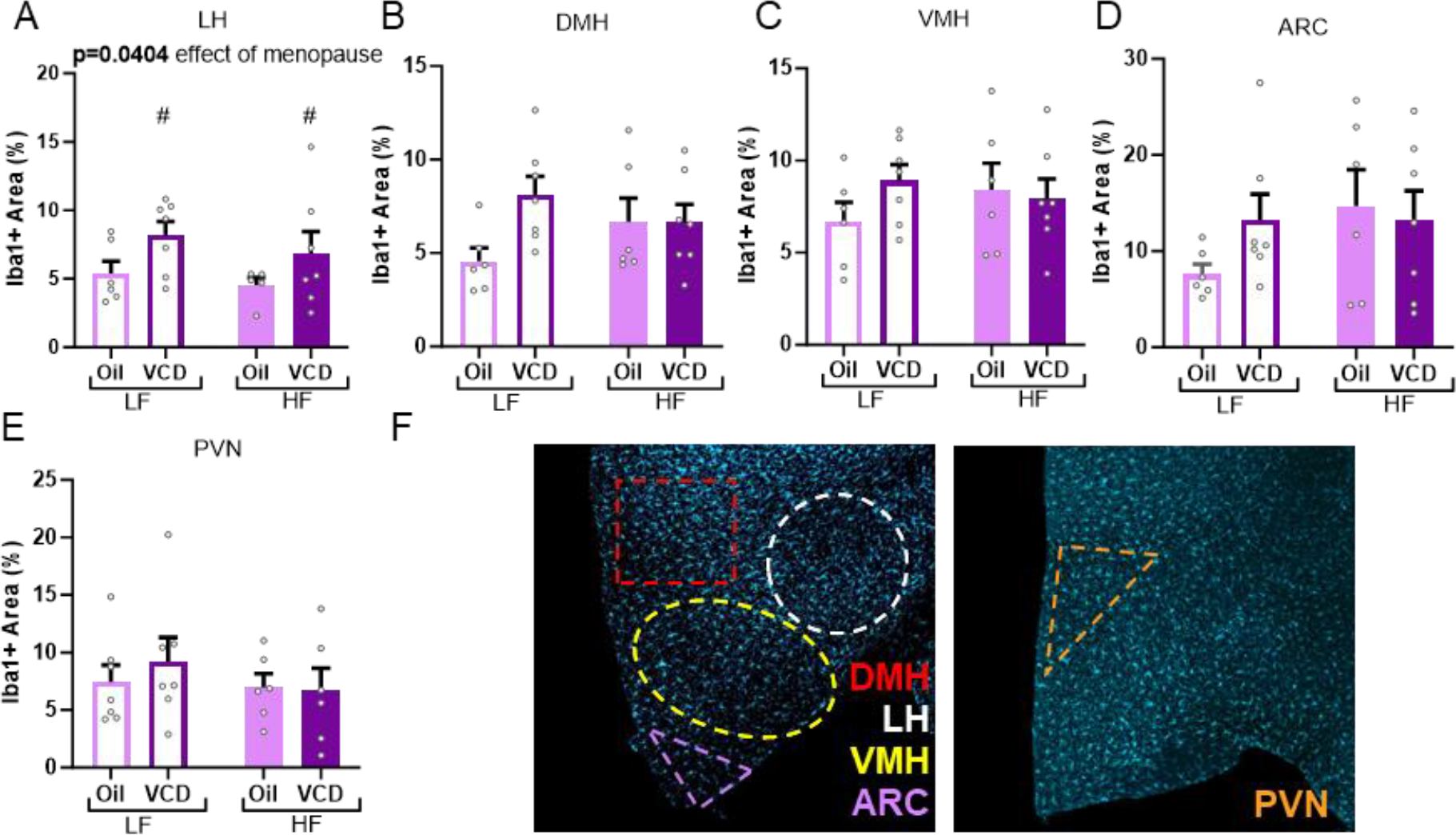
Interaction Between Menopause and HF Diet on Microglial Response in the Hypothalamus. Quantification of the area density (%) occupied by the Iba1 labeling in the (**A**) lateral hypothalamus area (LH), (**B**) dorsomedial hypothalamus (DMH), (**C**) ventromedial hypothalamus (VMH), (**D**) the arcuate nucleus (ARC), and (**E**) the paraventricular nucleus (PVN) of the hypothalamus. A 2way ANOVA was used to assess the effect of diet and menopause. (**F**) Representative images of microglia labeling using anti-Iba1 (Cyan, microglia marker). Dotted lines represent different regions of interest indicating different nuclei associated with the corresponding color. n=5-7 mice/group. High fat (HF), low fat (LF), 4-vinylcyclohexene diepoxide (VCD, menopause model).

### Menopause does not alter overall activity levels

General locomotor activity was assessed in the open field test. As expected, HF diet fed mice covered less distance; however, no effect of menopause was observed (**Sup Figure 2A**). Further, locomotor activity did not correlate with any of the metabolic measures (**Sup Figure 2B**), suggesting that changes in metabolic measures might not be mediated by physical activity.

## Discussion

AD patients often have metabolic dysfunction, which has been shown to be in part due to hypothalamic pathology (3, 7, 48). Female sex is also a risk factor for AD, such that ∼2/3 of AD patients are women (21, 22). Given that the majority of those who suffer from AD are elderly, most female AD patients are post-menopausal. On its own, menopause has been shown to increase metabolic risk factors for AD, such as obesity and diabetes (23, 49-52). Here, we sought to determine the effects of menopause on metabolic health in a mouse model of AD. Using an accelerated ovarian failure model in App^NL-F^ female mice, on either a LF or HF diet, menopausal mice exhibited worse outcomes on some metabolic measures. Our data further suggests that some of these effects of menopause may stem from central changes in the hypothalamus, such as altered expression of genes regulating energy balance and increased microgliosis.

Metabolic disease (including obesity and glucose intolerance) is a major dementia risk factor (4, 9-11). Further, glucose intolerance (diabetes and prediabetes) appears to confer an even greater risk for dementia in women compared to men (53, 54). Animal studies show that middle-aged female mice are more vulnerable to the negative metabolic effects of a high fat diet (36, 37). These findings also extend to mouse models of AD (5, 6, 55). For example, in both 3xTg-AD and TgAPP mice, females gain more weight, accumulate more visceral fat, and develop worse glucose intolerance than males or WT females (5, 55). Weight gain and glucose intolerance are known metabolic changes that can occur in women after menopause (49-52). These changes have also been described in the ovariectomy animal model of surgical menopause (56, 57) and in the VCD induced menopause model (35, 58, 59). In agreement with these studies, our study shows that, in the context of AD, menopausal female mice exhibit more severe obesity and worse glucose tolerance compared to controls.

To further characterize the metabolic state of these mice, we measured serum levels of diabetes-associated markers. We found that on a LF diet, VCD-injected menopausal females had significantly less serum insulin than oil-injected control females on a LF diet. This might explain the higher glucose levels observed in these mice at the time of GTT. Higher insulin levels were associated with increased body weight and worse glucose tolerance in VCD-injected menopausal mice. These findings indicated that the adverse metabolic effects observed in menopausal mice on a LF diet could be mediated by insulin dysregulation. Previous studies using this menopause model in WT mice do not report a difference in fasting insulin levels between control and VCD-injected mice on a control diet (58). However, on a HF diet, menopausal females had significantly higher fasting insulin levels than cycling controls (58). These differences could be due to the AD pathology in our mice that starts at around 6 months of age. Insulin resistance drastically increases in women after menopause (51, 60-62). Studies have shown that resistin levels vary in parallel and in the same direction as insulin and glucose levels (63, 64). In our study, menopause did not affect resistin levels. This is in line with other studies showing no change in resistin production by adipose tissue in ovariectomized female rats (65). This decoupling between insulin and resistin levels in LF-fed AD mice could be a potential mechanism behind the adverse effects of menopause on glucose tolerance. In this study, our samples were examined from non-fasting animals during their inactive period (light cycle); values for insulin and glucagon, among others, could be different if examined after a fasting period. Further studies, beyond the scope of this paper, are needed to elucidate the interaction between menopause and insulin signaling in the context of AD.

Balancing food consumption with energy expenditure is centrally regulated in the hypothalamus. This brain region is gaining interest in AD research as it has been shown to be affected by both amyloid plaques and tau neurofibrillary tangles, contributing to metabolic and non-cognitive deficits seen in AD patients that are observed prior to the onset of cognitive impairment (3, 7, 48). We therefore sought to explore how menopause and diet can influence changes in the hypothalamus in the context of AD. We found a decrease in Agrp expression in menopausal mice, specifically in those fed a LF diet. This effect could be due to the loss of estrogen. Estradiol is known to be anorexigenic and regulates food intake in part by inhibiting the excitability of the hypothalamic neuropeptide Y/agouti-related peptide (NPY/AgRP) neurons (66). Further, in ovariectomized female mice, treatment with estradiol restores the response of NPY/AgRP neurons to insulin (66). Aside from NPY/AgRP, estrogen has also been shown to promote POMC excitability (67); however, we did not observe any group differences in Pomc mRNA levels. Some of estrogen’s effects may also be mediated through peripheral effects on other organs, such as the liver. For example, conditional knock-out of estrogen receptor alpha in the liver leads to alterations of hepatic lipid metabolism and dysregulation of AgRP neurons (68). These results provide a mechanistic insight into the effects of estrogen loss induced by menopause in a mouse model of AD and suggest that AgRP may be a promising target for future studies.

AgRP also acts as an MC4R antagonist, driving energy intake (69). Menopausal mice tended to have higher Mc4r mRNA expression in the hypothalamus. MC4R is activated principally by α-melanocyte stimulating hormone (MSH, bioactive products of POMC processing), which promotes a cessation of feeding, increased energy expenditure and weight loss. Estrogen has been described to increase Mc4r gene expression via estrogen receptor alpha (70). Surprisingly, VCD mice tended to have higher Mc4r mRNA expression regardless of diet; however, studies have shown that androgens can also activate Mc4r gene in female rats (71, 72). Female rats treated with subcutaneous dihydrotestosterone pellets exhibit higher expression of Mc4r than controls (71). In women, postmenopausal ovaries stop producing estrogen but continue to produce androgens (73), a hormonal change that is also present in the VCD menopause model (74) but lacking in the ovariectomy model. The increase in Mc4r following menopause could be a compensatory mechanism, potentially induced by androgens, to reduce food consumption and prevent excessive weight gain following menopause. Further studies are needed to elucidate the link between androgens and Mc4r gene expression in postmenopausal mice. One limitation of the current study is that our findings are based on measurements of mRNA, which may not be translated on the protein level. Moreover, our findings are in dissections of the whole hypothalamus and do not allow for a regional analysis of changes in metabolic genes. Another limitation of this study is the lack of non-AD mice; however, in a previous study, we thoroughly compared wild type and 3xTg-AD mice and did not observe any effects of AD background on any of these hypothalamic markers in females (5).

A state of neuroinflammation, marked by microgliosis, has been described in both AD and metabolic disease (44-47). Studies show that hypothalamic inflammation has been linked to energy balance disruptions (3, 75-77). To test whether increased inflammation might contribute to the metabolic status of our mice, we assessed microgliosis in several hypothalamic nuclei involved in energy balance and glucose homeostasis. We found that menopause leads to increased microgliosis in the LH, independent of diet. The LH is linked to hunger and food intake. An increased microgliosis in menopausal female mice could be indicative of a disrupted state, potentially leading to adverse metabolic outcomes. In mice lacking estrogen receptor alpha signaling in the liver, dysregulation of AgRP neurons was associated with an increase state of microglia reactivity (68). These results indicate that increased hypothalamic inflammation might contribute to the deleterious metabolic effects of menopause.

Changes in hypothalamic gene expression and neuroinflammation observed above may result in changes in feeding behavior and/or energy expenditure. However, we do not have measurements of food intake, and/or indirect calorimetry to assess reasons underlying differences in metabolic outcomes. When testing general locomotor activity using the open field test, we did find that HF diet fed mice covered less distance, but no effect of menopause was observed. Further, locomotor activity did not correlate with any of the metabolic measures, suggesting that changes in metabolic measures might not be mediated by physical activity.

This is the first study to examine the interaction between metabolic disease modeled by a HF diet, and menopause using an accelerated ovarian failure model in an AD mouse model. This is an important contribution because the accelerated ovarian failure model is a follicle depleted ovary-intact model in which mice go through a peri-menopausal phase before entering a menopausal phase in which the ovaries cease to produce estrogens but continue to produce androgens (similar to post-menopausal women). We found that in addition to HF diet, menopause caused metabolic impairment in AD mice. Some effects of menopause may be centrally regulated by affecting hypothalamic expression of orexigenic genes and increasing inflammation in certain hypothalamic nuclei. How these metabolic changes affect AD pathology and cognitive impairment is the subject of other current studies in our lab. The current data show menopause-specific changes in metabolic control, highlighting the need to model menopause in preclinical studies of dementia (AD) and co-morbid risk factors such as metabolic disease.

## Authors’ Contributions

KLZ obtained funding for the experiments. CAG and KLZ designed the experiments with advisement from LSR, HLB and DGZ. CAG, AES, DR, RDK, DK, NK, OJG, MW, and FM performed the experiments. CAG, RDK, KAR, CAT, RMS, EG, KBM analyzed the data. CAG prepared the figures. CAG prepared the manuscript. KLZ and LSR edited the manuscript. All authors approved the final manuscript.

## Acknowledgements

The authors would like to thank Dr Takaomi Saido for providing the MTA for the App^tm2.1Tcs^/App^tm2.1Tcs^ (App^NL-F^ ) mice.

## Abbreviations

(VCD): 4-Vinylcyclohexene Diepoxide
(AD): Agouti-Related Peptide (AgRP) Alzheimer’s Disease
(ARC): Arcuate Nucleus
(AUC): Area Under the Curve
(DMH): Dorsomedial Hypothalamus
(GIP): Gastric Inhibitory Polypeptide
(GLP-1): Glucagon-Like-Peptide 1
(GTT): Glucose Tolerance Test
(HF): High Fat
(LH): Lateral Hypothalamus
(LepR): Leptin Receptor
(LF): Low Fat
(MC4R): Melanocortin 4 Receptor
(NPY): Neuropeptide Y
(PVN): Paraventricular Nucleus
(PAI-1): Plasminogen Activator Inhibitor-1
(POMC): Pro-Opiomelanocortin
(ROI): Regions Of Interest
(VMH): Ventromedial Hypothalamus

---

**Sup. Figure 1:**
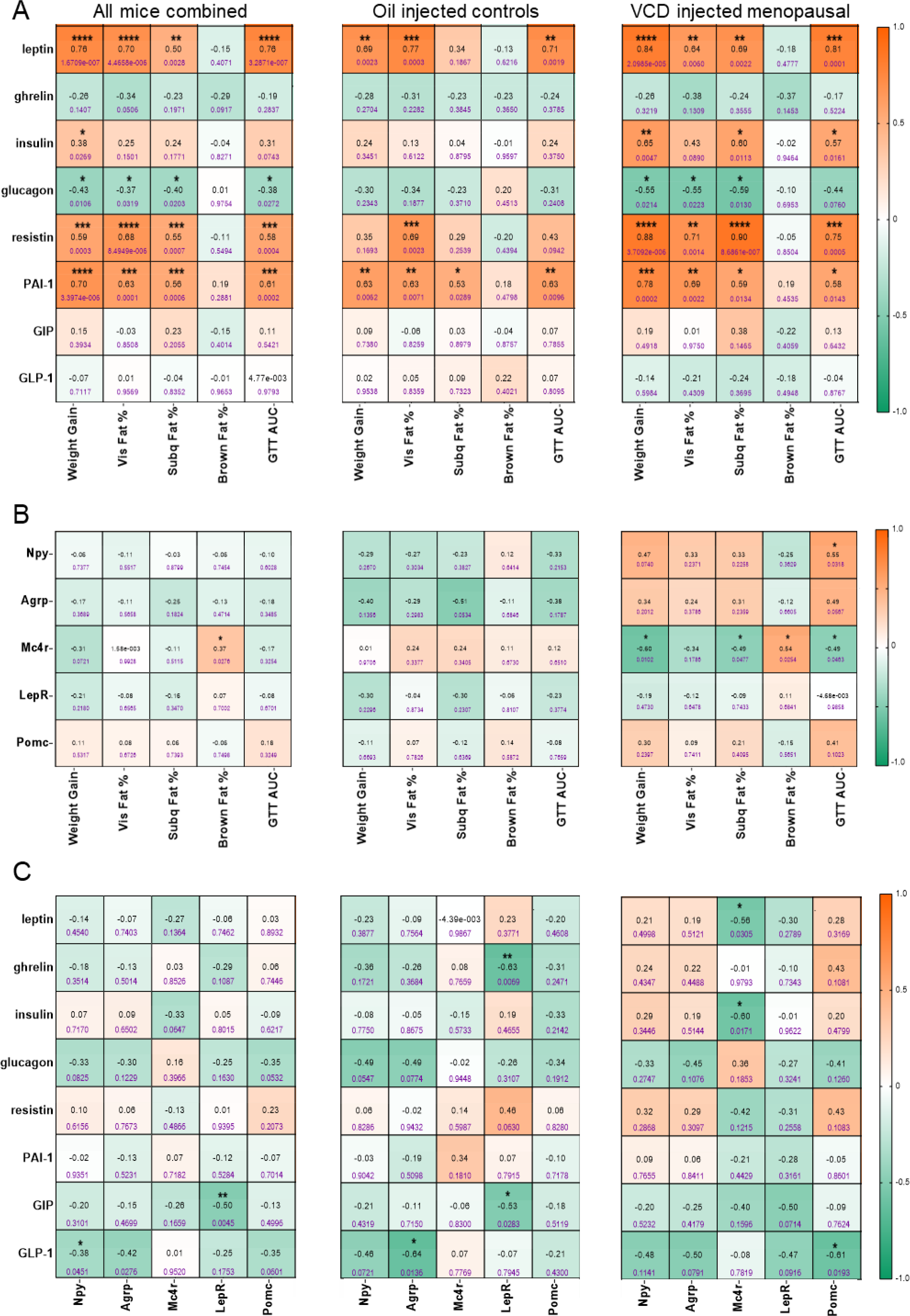
Correlations Between Different Metabolic Parameters. Pearson’s correlation matrices representing the relationships between (**A**) metabolic measures (body weight change; fat composition, and glucose intolerance), with serum levels of diabetes-associated markers; (**B**) metabolic measures with hypothalamic gene expression of energy expenditure related markers; (**C**) hypothalamic genes with serum markers. Left column tables are in all animals combined, middle column represents oil-injected control mice and right column shows correlations in VCD-injected menopausal mice. GTT AUC: area under the curve from the glucose tolerance test, high GTT AUC indicates greater glucose intolerance. Vis: visceral, Subq: subcutaneous, PAI-1: plasminogen activator inhibitor-1, GIP: gastric inhibitory polypeptide, GLP-1: glucagon-like-peptide 1. Pearson r values are presented in black, p-values are presented in purple font, *p<0.05, **p<0.01, ***p<0.001, ****p<0.0001 significant correlation; Orange: positive correlation, Green: negative correlation.

**Sup. Figure 2:**
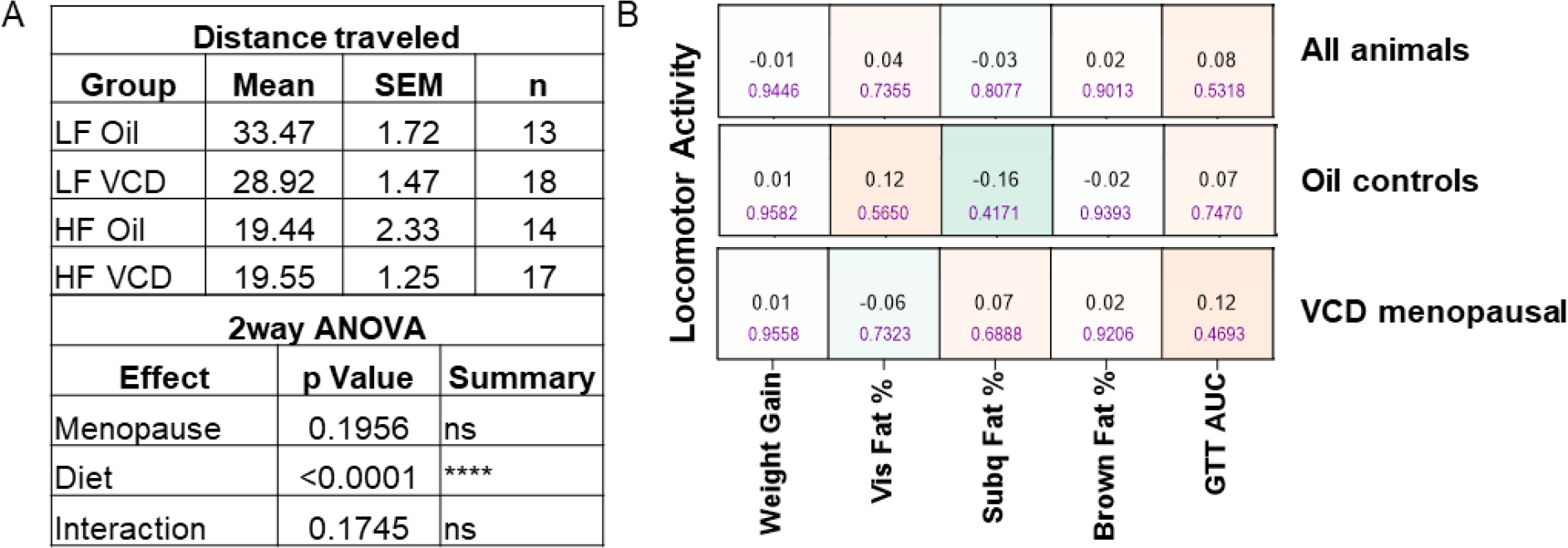
Locomotor Activity and Correlations With Metabolic Outcomes. (**A**) Table representing average and SEM distance traveled in an open field arena as a measure of locomotor activity as well as n numbers for each group. Results of the 2WAY ANOVA are described for each variable. (**B**) Pearson’s correlation matrices representing the relationships between metabolic measures (body weight change; fat composition, and glucose intolerance), with locomotor activity. Pearson’s r values are presented in black; p-values are presented in purple font. Orange: positive correlation, Green: negative correlation. GTT AUC: area under the curve from the glucose tolerance test, high GTT AUC indicates greater glucose intolerance. VCD: 4-vinylcyclohexene diepoxide (menopause model). Vis: visceral, Subq: subcutaneous.

## References

1. Gustavsson A, Norton N, Fast T, Frolich L, Georges J, Holzapfel D, et al. Global estimates on the number of persons across the Alzheimer’s disease continuum. Alzheimers Dement. 2023;19(2):658–70.

2. Matthews KA, Xu W, Gaglioti AH, Holt JB, Croft JB, Mack D, et al. Racial and ethnic estimates of Alzheimer’s disease and related dementias in the United States (2015-2060) in adults aged >/=65 years. Alzheimers Dement. 2019;15(1):17–24.

3. Ishii M, Iadecola C. Metabolic and Non-Cognitive Manifestations of Alzheimer’s Disease: The Hypothalamus as Both Culprit and Target of Pathology. Cell Metab. 2015;22(5):761–76.

4. Onaolapo AY, Ojo FO, Adeleye OO, Falade J, Onaolapo OJ. Diabetes mellitus and energy dysmetabolism in Alzheimer’s disease: Understanding the relationships and potential therapeutic targets. Curr Diabetes Rev. 2023.

5. Robison LS, Gannon OJ, Thomas MA, Salinero AE, Abi-Ghanem C, Poitelon Y, et al. Role of sex and high-fat diet in metabolic and hypothalamic disturbances in the 3xTg-AD mouse model of Alzheimer’s disease. Journal of neuroinflammation. 2020;17(1):285.

6. Robison LS, Gannon OJ, Salinero AE, Abi-Ghanem C, Kelly RD, Riccio DA, et al. Sex differences in metabolic phenotype and hypothalamic inflammation in the 3xTg-AD mouse model of Alzheimer’s disease. Biol Sex Differ. 2023;14(1):51.

7. Ishii M, Wang G, Racchumi G, Dyke JP, Iadecola C. Transgenic mice overexpressing amyloid precursor protein exhibit early metabolic deficits and a pathologically low leptin state associated with hypothalamic dysfunction in arcuate neuropeptide Y neurons. J Neurosci. 2014;34(27):9096–106.

8. Zheng H, Zhou Q, Du Y, Li C, Xu P, Lin L, et al. The hypothalamus as the primary brain region of metabolic abnormalities in APP/PS1 transgenic mouse model of Alzheimer’s disease. Biochim Biophys Acta Mol Basis Dis. 2018;1864(1):263–73.

9. Xue M, Xu W, Ou YN, Cao XP, Tan MS, Tan L, et al. Diabetes mellitus and risks of cognitive impairment and dementia: A systematic review and meta-analysis of 144 prospective studies. Ageing research reviews. 2019;55:100944.

10. Whitmer RA, Gunderson EP, Quesenberry CP, Jr., Zhou J, Yaffe K. Body mass index in midlife and risk of Alzheimer disease and vascular dementia. Current Alzheimer research. 2007;4(2):103–9.

11. Kim YJ, Kim SM, Jeong DH, Lee SK, Ahn ME, Ryu OH. Associations between metabolic syndrome and type of dementia: analysis based on the National Health Insurance Service database of Gangwon province in South Korea. Diabetol Metab Syndr. 2021;13(1).

12. Janson J, Laedtke T, Parisi JE, O’Brien P, Petersen RC, Butler PC. Increased risk of type 2 diabetes in Alzheimer disease. Diabetes. 2004;53(2):474–81.

13. Moloney AM, Griffin RJ, Timmons S, O’Connor R, Ravid R, O’Neill C. Defects in IGF-1 receptor, insulin receptor and IRS-1/2 in Alzheimer’s disease indicate possible resistance to IGF-1 and insulin signalling. Neurobiol Aging. 2010;31(2):224–43.

14. Talbot K, Wang HY, Kazi H, Han LY, Bakshi KP, Stucky A, et al. Demonstrated brain insulin resistance in Alzheimer’s disease patients is associated with IGF-1 resistance, IRS-1 dysregulation, and cognitive decline. J Clin Invest. 2012;122(4):1316–38.

15. Ho L, Qin W, Pompl PN, Xiang Z, Wang J, Zhao Z, et al. Diet-induced insulin resistance promotes amyloidosis in a transgenic mouse model of Alzheimer’s disease. FASEB J. 2004;18(7):902–4.

16. Julien C, Tremblay C, Phivilay A, Berthiaume L, Emond V, Julien P, et al. High-fat diet aggravates amyloid-beta and tau pathologies in the 3xTg-AD mouse model. Neurobiol Aging. 2010;31(9):1516–31.

17. Barron AM, Rosario ER, Elteriefi R, Pike CJ. Sex-specific effects of high fat diet on indices of metabolic syndrome in 3xTg-AD mice: implications for Alzheimer’s disease. PloS one. 2013;8(10):e78554.

18. Bracko O, Vinarcsik LK, Cruz Hernández JC, Ruiz-Uribe NE, Haft-Javaherian M, Falkenhain K, et al. High fat diet worsens Alzheimer’s disease-related behavioral abnormalities and neuropathology in APP/PS1 mice, but not by synergistically decreasing cerebral blood flow. Scientific reports. 2020;10(1):9884.

19. Takeda S, Sato N, Uchio-Yamada K, Sawada K, Kunieda T, Takeuchi D, et al. Diabetes-accelerated memory dysfunction via cerebrovascular inflammation and Abeta deposition in an Alzheimer mouse model with diabetes. Proc Natl Acad Sci U S A. 2010;107(15):7036–41.

20. Gannon OJ, Robison LS, Salinero AE, Abi-Ghanem C, Mansour FM, Kelly RD, et al. Highfat diet exacerbates cognitive decline in mouse models of Alzheimer’s disease and mixed dementia in a sex-dependent manner. Journal of neuroinflammation. 2022;19(1):110.

21. 2020 Alzheimer’s disease facts and figures. Alzheimers Dement. 2020.

22. 2023 Alzheimer’s disease facts and figures. Alzheimers Dement. 2023;19(4):1598–695.

23. El Khoudary SR, Aggarwal B, Beckie TM, Hodis HN, Johnson AE, Langer RD, et al. Menopause Transition and Cardiovascular Disease Risk: Implications for Timing of Early Prevention: A Scientific Statement From the American Heart Association. Circulation. 2020;142(25):e506–e32.

24. Jeong HG, Park H. Metabolic Disorders in Menopause. Metabolites. 2022;12(10).

25. Lisabeth L, Bushnell C. Stroke risk in women: the role of menopause and hormone therapy. Lancet Neurol. 2012;11(1):82–91.

26. Rocca WA, Grossardt BR, Maraganore DM. The long-term effects of oophorectomy on cognitive and motor aging are age dependent. Neurodegener Dis. 2008;5(3-4):257–60.

27. Stachowiak G, Pertynski T, Pertynska-Marczewska M. Metabolic disorders in menopause. Prz Menopauzalny. 2015;14(1):59–64.

28. El Khoudary SR, Aggarwal B, Beckie TM, Hodis HN, Johnson AE, Langer RD, et al. Menopause Transition and Cardiovascular Disease Risk: Implications for Timing of Early Prevention: A Scientific Statement From the American Heart Association. Circulation. 2020;142(25):e506–e32.

29. Cheng G, Huang C, Deng H, Wang H. Diabetes as a risk factor for dementia and mild cognitive impairment: a meta-analysis of longitudinal studies. Intern Med J. 2012;42(5):484–91.

30. Pirimoglu ZM, Arslan C, Buyukbayrak EE, Kars B, Karsidag YK, Unal O, et al. Glucose tolerance of premenopausal women after menopause due to surgical removal of ovaries. Climacteric. 2011;14(4):453–7.

31. Slopien R, Wender-Ozegowska E, Rogowicz-Frontczak A, Meczekalski B, Zozulinska-Ziolkiewicz D, Jaremek JD, et al. Menopause and diabetes: EMAS clinical guide. Maturitas. 2018;117:6–10.

32. Mosconi L, Rahman A, Diaz I, Wu X, Scheyer O, Hristov HW, et al. Increased Alzheimer’s risk during the menopause transition: A 3-year longitudinal brain imaging study. PLoS One. 2018;13(12):e0207885.

33. Saito T, Matsuba Y, Mihira N, Takano J, Nilsson P, Itohara S, et al. Single App knock-in mouse models of Alzheimer’s disease. Nature neuroscience. 2014;17(5):661–3.

34. Brooks HL, Pollow DP, Hoyer PB. The VCD Mouse Model of Menopause and Perimenopause for the Study of Sex Differences in Cardiovascular Disease and the Metabolic Syndrome. Physiology (Bethesda). 2016;31(4):250–7.

35. Gannon OJ, Naik JS, Riccio D, Mansour FM, Abi-Ghanem C, Salinero AE, et al. Menopause causes metabolic and cognitive impairments in a chronic cerebral hypoperfusion model of vascular contributions to cognitive impairment and dementia. Biology of sex differences. 2023;14(1):34.

36. Salinero AE, Robison LS, Gannon OJ, Riccio D, Mansour F, Abi-Ghanem C, et al. Sexspecific effects of high-fat diet on cognitive impairment in a mouse model of VCID. FASEB journal: official publication of the Federation of American Societies for Experimental Biology. 2020;34(11):15108–22.

37. Salinero AE, Anderson BM, Zuloaga KL. Sex differences in the metabolic effects of diet-induced obesity vary by age of onset. Int J Obes (Lond). 2018;42(5):1088–91.

38. Robison LS, Albert NM, Camargo LA, Anderson BM, Salinero AE, Riccio DA, et al. High-Fat Diet-Induced Obesity Causes Sex-Specific Deficits in Adult Hippocampal Neurogenesis in Mice. eNeuro. 2020;7(1).

39. Klok MD, Jakobsdottir S, Drent ML. The role of leptin and ghrelin in the regulation of food intake and body weight in humans: a review. Obes Rev. 2007;8(1):21–34.

40. Cesari M, Pahor M, Incalzi RA. Plasminogen activator inhibitor-1 (PAI-1): a key factor linking fibrinolysis and age-related subclinical and clinical conditions. Cardiovasc Ther. 2010;28(5):e72–91.

41. Seino Y, Fukushima M, Yabe D. GIP and GLP-1, the two incretin hormones: Similarities and differences. J Diabetes Investig. 2010;1(1-2):8–23.

42. Farooqi IS, Keogh JM, Yeo GS, Lank EJ, Cheetham T, O’Rahilly S. Clinical spectrum of obesity and mutations in the melanocortin 4 receptor gene. N Engl J Med. 2003;348(12):1085–95.

43. Huszar D, Lynch CA, Fairchild-Huntress V, Dunmore JH, Fang Q, Berkemeier LR, et al. Targeted disruption of the melanocortin-4 receptor results in obesity in mice. Cell. 1997;88(1):131–41.

44. Hansen DV, Hanson JE, Sheng M. Microglia in Alzheimer’s disease. J Cell Biol. 2018;217(2):459–72.

45. Baufeld C, Osterloh A, Prokop S, Miller KR, Heppner FL. High-fat diet-induced brain region-specific phenotypic spectrum of CNS resident microglia. Acta Neuropathol. 2016;132(3):361–75.

46. Thaler JP, Yi CX, Schur EA, Guyenet SJ, Hwang BH, Dietrich MO, et al. Obesity is associated with hypothalamic injury in rodents and humans. J Clin Invest. 2012;122(1):153–62.

47. Maldonado-Ruiz R, Montalvo-Martinez L, Fuentes-Mera L, Camacho A. Microglia activation due to obesity programs metabolic failure leading to type two diabetes. Nutr Diabetes. 2017;7(3):e254.

48. Braak H, Braak E. Neuropathological stageing of Alzheimer-related changes. Acta Neuropathol. 1991;82(4):239–59.

49. Otsuki M, Kasayama S, Morita S, Asanuma N, Saito H, Mukai M, et al. Menopause, but not age, is an independent risk factor for fasting plasma glucose levels in nondiabetic women. Menopause. 2007;14(3 Pt 1):404–7.

50. Wu SI, Chou P, Tsai ST. The impact of years since menopause on the development of impaired glucose tolerance. J Clin Epidemiol. 2001;54(2):117–20.

51. Phillips GB, Jing T, Heymsfield SB. Does insulin resistance, visceral adiposity, or a sex hormone alteration underlie the metabolic syndrome? Studies in women. Metabolism. 2008;57(6):838–44.

52. Tremollieres FA, Pouilles JM, Ribot CA. Relative influence of age and menopause on total and regional body composition changes in postmenopausal women. Am J Obstet Gynecol. 1996;175(6):1594–600.

53. Chatterjee S, Peters SA, Woodward M, Mejia Arango S, Batty GD, Beckett N, et al. Type 2 Diabetes as a Risk Factor for Dementia in Women Compared With Men: A Pooled Analysis of 2.3 Million People Comprising More Than 100,000 Cases of Dementia. Diabetes Care. 2016;39(2):300–7.

54. Sundermann EE, Thomas KR, Bangen KJ, Weigand AJ, Eppig JS, Edmonds EC, et al. Prediabetes Is Associated With Brain Hypometabolism and Cognitive Decline in a Sex-Dependent Manner: A Longitudinal Study of Nondemented Older Adults. Frontiers in neurology. 2021;12:551975.

55. Freire-Regatillo A, Diaz-Pacheco S, Frago LM, Arevalo MA, Argente J, Garcia-Segura LM, et al. Sex Differences in Hypothalamic Changes and the Metabolic Response of TgAPP Mice to a High Fat Diet. Front Neuroanat. 2022;16:910477.

56. Rogers NH, Perfield JW, 2nd, Strissel KJ, Obin MS, Greenberg AS. Reduced energy expenditure and increased inflammation are early events in the development of ovariectomy-induced obesity. Endocrinology. 2009;150(5):2161–8.

57. Stokar J, Gurt I, Cohen-Kfir E, Yakubovsky O, Hallak N, Benyamini H, et al. Hepatic adropin is regulated by estrogen and contributes to adverse metabolic phenotypes in ovariectomized mice. Mol Metab. 2022;60:101482.

58. Romero-Aleshire MJ, Diamond-Stanic MK, Hasty AH, Hoyer PB, Brooks HL. Loss of ovarian function in the VCD mouse-model of menopause leads to insulin resistance and a rapid progression into the metabolic syndrome. Am J Physiol Regul Integr Comp Physiol. 2009;297(3):R587–92.

59. Sui K, Yasrebi A, Longoria CR, MacDonell AT, Jaffri ZH, Martinez SA, et al. Coconut Oil Saturated Fatty Acids Improved Energy Homeostasis but not Blood Pressure or Cognition in VCD-Treated Female Mice. Endocrinology. 2023;164(3).

60. C VS, S B, A S. Analysis of the degree of insulin resistance in post menopausal women by using skin temperature measurements and fasting insulin and fasting glucose levels: a case control study. J Clin Diagn Res. 2012;6(10):1644–7.

61. Cho GJ, Lee JH, Park HT, Shin JH, Hong SC, Kim T, et al. Postmenopausal status according to years since menopause as an independent risk factor for the metabolic syndrome. Menopause. 2008;15(3):524–9.

62. Innes KE, Selfe TK, Taylor AG. Menopause, the metabolic syndrome, and mind-body therapies. Menopause. 2008;15(5):1005–13.

63. Abdalla MMI. Salivary resistin level and its association with insulin resistance in obese individuals. World J Diabetes. 2021;12(9):1507–17.

64. Rajala MW, Qi Y, Patel HR, Takahashi N, Banerjee R, Pajvani UB, et al. Regulation of resistin expression and circulating levels in obesity, diabetes, and fasting. Diabetes. 2004;53(7):1671–9.

65. Nogueiras R, Gualillo O, Caminos JE, Casanueva FF, Dieguez C. Regulation of resistin by gonadal, thyroid hormone, and nutritional status. Obes Res. 2003;11(3):408–14.

66. Qiu J, Bosch MA, Zhang C, Ronnekleiv OK, Kelly MJ. Estradiol Protects Neuropeptide Y/Agouti-Related Peptide Neurons against Insulin Resistance in Females. Neuroendocrinology. 2020;110(1-2):105–18.

67. Stincic TL, Grachev P, Bosch MA, Ronnekleiv OK, Kelly MJ. Estradiol Drives the Anorexigenic Activity of Proopiomelanocortin Neurons in Female Mice. eNeuro. 2018;5(4).

68. Benedusi V, Della Torre S, Mitro N, Caruso D, Oberto A, Tronel C, et al. Liver ERalpha regulates AgRP neuronal activity in the arcuate nucleus of female mice. Sci Rep. 2017;7(1):1194.

69. Krashes MJ, Lowell BB, Garfield AS. Melanocortin-4 receptor-regulated energy homeostasis. Nat Neurosci. 2016;19(2):206–19.

70. Krause WC, Rodriguez R, Gegenhuber B, Matharu N, Rodriguez AN, Padilla-Roger AM, et al. Oestrogen engages brain MC4R signalling to drive physical activity in female mice. Nature. 2021;599(7883):131–5.

71. Maranon R, Lima R, Spradley FT, do Carmo JM, Zhang H, Smith AD, et al. Roles for the sympathetic nervous system, renal nerves, and CNS melanocortin-4 receptor in the elevated blood pressure in hyperandrogenemic female rats. Am J Physiol Regul Integr Comp Physiol. 2015;308(8):R708–13.

72. Reckelhoff JF. Androgens and Blood Pressure Control: Sex Differences and Mechanisms. Mayo Clin Proc. 2019;94(3):536–43.

73. Brzozowska M, Lewinski A. Changes of androgens levels in menopausal women. Prz Menopauzalny. 2020;19(4):151–4.

74. Mayer LP, Devine PJ, Dyer CA, Hoyer PB. The follicle-deplete mouse ovary produces androgen. Biol Reprod. 2004;71(1):130–8.

75. Arruda AP, Milanski M, Coope A, Torsoni AS, Ropelle E, Carvalho DP, et al. Low-grade hypothalamic inflammation leads to defective thermogenesis, insulin resistance, and impaired insulin secretion. Endocrinology. 2011;152(4):1314–26.

76. Wisse BE, Ogimoto K, Tang J, Harris MK, Jr., Raines EW, Schwartz MW. Evidence that lipopolysaccharide-induced anorexia depends upon central, rather than peripheral, inflammatory signals. Endocrinology. 2007;148(11):5230–7.

77. Le Thuc O, Stobbe K, Cansell C, Nahon JL, Blondeau N, Rovere C. Hypothalamic Inflammation and Energy Balance Disruptions: Spotlight on Chemokines. Front Endocrinol (Lausanne). 2017;8:197.

